# Conditional independence as a statistical assessment of evidence integration processes

**DOI:** 10.1101/2023.05.03.539321

**Authors:** Emilio Salinas, Terrence R Stanford

## Abstract

Intuitively, combining multiple sources of evidence should lead to more accurate decisions than considering single sources of evidence individually. In practice, however, the proper computation may be difficult, or may require additional data that are inaccessible. Here, based on the concept of conditional independence, we consider expressions that can serve either as recipes for integrating evidence based on limited data, or as statistical benchmarks for characterizing evidence integration processes. Consider three events, *A, B*, and *C*. We find that, if *A* and *B* are conditionally independent with respect to *C*, then the probability that *C* occurs given that both *A* and *B* are known, *P* (*C*|*A, B*), can be easily calculated without the need to measure the full three-way dependency between *A, B*, and *C*. This simplified approach can be used in two general ways: to generate predictions by combining multiple (conditionally independent) sources of evidence, or to test whether separate sources of evidence are functionally independent of each other. These applications are demonstrated with four computer-simulated examples, which include detecting a disease based on repeated diagnostic testing, inferring biological age based on multiple biomarkers of aging, discriminating two spatial locations based on multiple cue stimuli (multisensory integration), and examining how behavioral performance in a visual search task depends on selection histories. Besides providing a sound prescription for predicting outcomes, this methodology may be useful for analyzing experimental data of many types.

## Introduction

Decisions are often based on multiple sources of evidence. For instance, to determine whether a patient has an infection, a doctor may take their temperature and order a lab test, and combine both results to form a diagnosis. Two interrelated questions about this process are key. One is how much information these two pieces of evidence provide individually versus together, and the other is how exactly they should be combined. Here we show that a relationship known as conditional independence can be used as a statistical tool and applied to data to help answer these questions.

The decision-making process requires two ingredients: computational recipes and knowledge of past outcomes under equivalent circumstances. The equivalence is key; for accurate estimation, the prior data must conform to the specific evidence samples being evaluated. In the case of the doctor’s diagnosis, the patient’s results should be compared (via appropriate probabilistic calculus) to prior measurements of temperature, lab test data, and infection status *taken simultaneously*. That is, in general, knowing the relationship between temperature and infection status on one hand, and lab test and infection status on the other, separately, is not sufficient for making an optimal informed decision — unless, as shown below, a particular relationship is satisfied between the three variables. In that case, optimal decisions and inferences can indeed be made based on more limited data.

This particular relationship is conditional independence [1]. When two sources of evidence, *A* and *B*, are conditionally independent with respect to a third quantity, *C*, then characterizing separately the relationships between *A* and *C* and between *B* and *C* suffices for completely characterizing how *C* will vary when *A* and *B* are both known; there is no need to make prior, simultaneous, three-way measurements to establish the necessary reference. This is important because, in practice, making pairwise measurements generally requires many fewer resources than making simultaneous measurements of more than two variables, everything else being equal, and the difference becomes dramatically larger as the number of variables increases beyond three.

Here we derive expressions for the performance that is expected when two or more sources of evidence are conditionally independent with respect to a variable of interest, and demonstrate their applicability to the life sciences with four examples. These expressions are potentially useful in two broad types of situation. One is when there are good reasons to *assume* that the conditional independence requirement is satisfied. In that case, the results can be used to make predictions easily; that is, to establish expected outcomes based on multiple, current evidence samples together with prior knowledge of pairwise measurements only. The other scenario in which the derived expressions are potentially useful is when simultaneous measurements of all the variables of interest are available. In that case, conditional independence, as construed here, serves as an intuitive benchmark to *test* whether separate pieces of evidence are influencing an outcome independently or not, which can provide insight into the underlying mechanistic interactions giving rise to the data. The concrete examples are chosen for clarity and to be representative of real-life contexts in which these situations may arise, but the range of potential applications is wide.

## Methods

All numerical calculations were performed using the Matlab programming environment (version R2013b and later; The Mathworks, Natick, MA). All data, analysis files, and code for generating the shown examples and plots are available from the Zenodo repository at https://doi.org/10.5281/zenodo.7894052. Functions for computing probabilities according to Equations 16, 37–38 are included as part of the software package.

### Calculating uncertainties in probability

The Results section presents expressions for estimating the probabilities *P* (*C*|*A, B*) based on the conditional probabilities *P* (*C*|*A*) and *P* (*C*|*B*) and the prior *P* (*C*). Uncertainties for the estimated probabilities were generated through a resampling procedure [2, 3]. The procedure requires knowing the numbers of observations associated with the measurement of each of the input probabilities, *P* (*C*|*A*), *P* (*C*|*B*), and *P* (*C*). With the numbers of observations at hand, it makes use of binomial statistics to generate distributions for the respective input probabilities. These distributions are used to recompute the estimated probability *P* (*C*|*A, B*) multiple times to generate a distribution for it, which can then be used to extract significance values or measures of dispersion. The Matlab code mentioned in the previous paragraph produces confidence intervals using this method. Example confidence intervals are shown in Fig. 6d.

This method is appropriate for evaluating the uncertainty associated with estimated individual probability terms at given values of *A, B*, and *C*. Separate statistical methods exist for evaluating whether three variables satisfy conditional independence [4, 5].

### Generating correlated outcomes

Simulating the outcome of a diagnostic test, a coin flip, or any other binary outcome is simple; it amounts to drawing a random sample from a uniform distribution and comparing it to the event’s probability [6]. To generate positively correlated data for *N* events (i.e., *N* diagnostic tests or biomarker readings), we used two methods. Both methods start by drawing a set of *N* independent random samples, *u*_*j*_, (with *j* = 1, …, *N*), and both use a parameter *α* that varies between 0 and 1 to determine the level of correlation between them. In method 1, an additional random sample *η* is drawn, and each of the *N* samples is reset to equal *η* with a probability *α*. The correlated random sample *r*_*j*_ is then

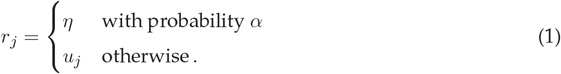

Thus, as *α* increases, it becomes more likely that a subset of the random samples are identical to each other, and thus that the outcomes of the simulated events are the same. Method 2 also starts with *N* independent random samples, *u*_*j*_, and an additional one, *η*, but now the *N* random samples move toward *η* by an amount that is proportional to *α*. In this case,

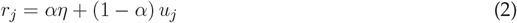

so *α* determines how close the correlated samples are to each other; when *α* = 1, they all converge on the same point.

Data shown in the figures are based on method 1, but results were qualitatively the same with method 2. The Matlab code that implements these methods is included in the deposited software package.

### Simulating behavior in the oddball task

The model for the oddball task simulates the performance of a subject. It takes a random sequence of target locations (with values 1, 2, 3, 4) and a random sequence of target colors (with values 1, 2) as inputs and produces a sequence of choices (also within locations 1, 2, 3, 4) as output. Each choice may be correct, if it matches the corresponding target location, or incorrect, if it does not. In each simulated trial, the choice is either a guess or an informed decision. The probability of making an informed decision is given by

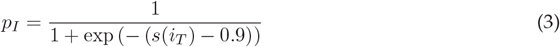

where *i*_*T*_ is the index of the target color (1 or 2) on the given trial, and *s*(*i*_*T*_) is the sensitivity associated with that color. Thus, the higher this sensitivity, the higher the probability of making an informed decision. A random number is drawn and compared to *p*_*I*_ to determine if the decision is informed or not, and if it is, the choice has a 0.95 probability of being correct. If the decision is not informed, then it is a guess. In that case, there are four numbers (≥ 0) to consider, *b*(1), *b*(2), *b*(3) and *b*(4), which represent the internal biases associated with each of the four locations. The bias for a given location is proportional to the probability that a guess is made toward that location. So, to generate a guess, a random number is drawn and compared to the four biases to determine the location corresponding to the choice. For example, if the biases have values [0 1 2 2], the probabilities associated with each location are [0 0.2 0.4 0.4], which add up to 1; then, to make a guess, we first compute the cumulative sum of the probability vector, [0 0.2 0.6 1] in this case, and then draw a random number between 0 and 1 from a uniform distribution and compare it to the cumulative values. In the example, if the number is between 0 and 0.2, the guess is toward location 2; if the number is between 0.2 and 0.6, the guess is toward location 3; and if the number is between 0.6 and 1, the guess is toward location 4. Finally, if the guessed location matches the target location, the choice is correct, otherwise it is an error.

The two color sensitivites and four location biases are updated after each choice, and in both cases the update consists of two parts. For the motor biases, the update rules are

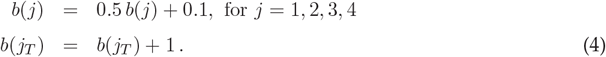

The first step tends to equalize the four biases, so that future guesses are more even across locations; the second step increases the bias associated with the target location, *b*(*j*_*T*_), so that future guesses toward it become more likely. These rules apply identically after all choices, whether correct or incorrect. For the color sensitivities the update rules are

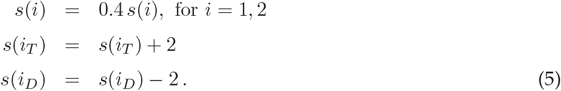

The first step moves the two sensitivies toward zero, making them more similar to each other; the second and third steps make them more different by increasing the sensistivity associated with the color of the target, *s*(*i*_*T*_), and decreasing the sensitivity associated with the color of the distracters, *s*(*i*_*D*_). Importantly, the first step applies after all choices whereas the other two apply only after correct choices. Note that, as per Equation 3, the larger the difference between *s*(*i*_*T*_) and *s*(*i*_*D*_), the larger the difference in performance between a target of one color or the other. Each simulation of the task ran for 100,000 trials.

For analysis of the simulated task data, the target color and target location in a series of consecutive trials can be encoded literally, as in {Red, Green, Red, Red}, or as same/different relative to the oddball color in the current trial, as in {Same, Different, Same, Same}. We use the latter encoding because it automatically pools the data for red and green targets, which were statistically identical. Likewise for target location; we use the same/different encoding because all four locations were statistically identical. Note, however, that in this case there are three possible non-target locations in each trial, and all of them count as different from the target regardless of whether they are repeated.

## Results and Discussion

### Predicting outcomes based on conditionally independent sources of evidence

The starting concept is that of conditional independence. Two events *A* and *B*, are said to be conditionally independent relative to a third event *C* if their joint probability given *C* is

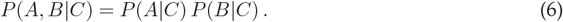

This means that, if *C* is known, *A* and *B* occur independently of each other. Said differently, if condition *C* is known, the probability that *B* occurs is fixed, regardless of whether *A* occurs or not. Indeed, the equality

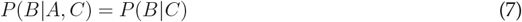

which directly states this idea, is equivalent to the first expression. Conditional independence is entirely separate from general independence; that is, Equation 6 does not tell us whether

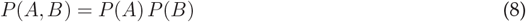

is true or not, and this relationship has no bearing on the results that follow. This is important.

Here is a concrete example presented as a frequency tree (Fig 1), which facilitates probabilistic reasoning and calculation [7, 8]. The top of the tree divides the day into two periods, before 6 pm (*A* = 0) and after 6 pm (*A* = 1) with equal probabilities (1/2). Now suppose that the probability that Carol is home (*C* = 1) is 0.6 when it is after 6 pm, but only 0.2 when it is before 6 pm, and furthermore, her dog only barks (*B* = 1) with a probability of 0.1 when Carol is home, but barks with a probability of 0.6 when she is away (*C* = 0). Note that the barking depends only on whether Carol is home or not, so knowing the time of day makes no difference in assessing what the dog will do, if we already know where Carol is. Translated into numbers, this means that

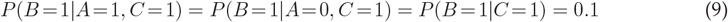

which can be easily verified from Fig 1. For instance, to calculate *P* (*B* = 1|*A* = 0, *C* = 1), count the number of occurrences where the three events are in the required states (bottom branch number 4 from the left) and divide by the number of occurrences where the two conditioning events, *A* and *C*, are in the required states, *A* = 0 and *C* = 1 (sum of bottom branches 3 and 4); this gives 1*/*10. Similarly, for *P* (*B* = 1|*C* = 1), the number of occurrences where *B* = 1 and *C* = 1 divided by the number where *C* = 1 gives 4*/*40. Equalities analogous to those in Equation 9 are also satisfied

**Fig 1.**
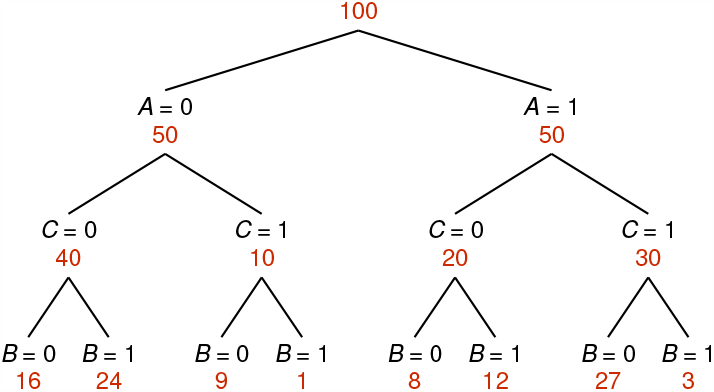
Frequency tree for three events. *A* corresponds to ‘time is after 6 pm’, *C* corresponds to ‘Carol is home’, and *B* corresponds to ‘the dog is barking’, with 1 and 0 indicating true and false. Numbers of observations are in red. Probabilities for all event combinations can be readily calculated from the tree. For example, the joint probability that all three events occur simultaneoulsy, *P* (*A* = 1, *B* = 1, *C* = 1), is equal to 3/100, as given by the rightmost branch. The probability that Carol is not home given that time is not past 6 pm, *P* (*C* = 0|*A* = 0), is equal to 40/50, as given by the leftmost branch, and so on. This example is constructed so that *A* and *B* occur independently of each other given *C*.

when *C* = 0 and for the case *B* = 0, in agreement with conditional independence as defined in Equation 7. However, *in general* the barking is not independent of time; for instance

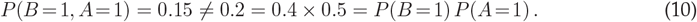

The crucial observation about this example is that, because of their conditional independence, the dog barking and the time of day can be used as independent pieces of evidence to estimate whether Carol is home or not. Now the quantity that we seek is the conditional probability of *C* given *A* and *B*, or

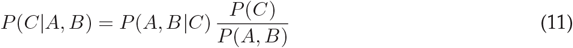

where the right side is the standard identity based on Bayes’ theorem. The calculation has no general solution because there is no universal way to obtain the first term on the right; the three-way relationship between *A, B*, and *C* simply has to be measured — unless *A* and *B* are conditionally independent given *C*. If that is the case, then Equation 6 can be used to factorize all the terms where *A* and *B* appear jointly in Equation 11, and everything can be put in terms of *P* (*C*|*A*) and *P* (*C*|*B*), which are the separate, individual pieces of information about the event of interest, whether Carol is home or not.

To derive the result, the first step is to rewrite the conditional independence constraint as

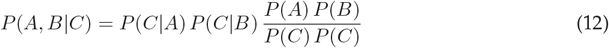

by applying Bayes’ rule to the two conditional probabilities in Equation 6. Next, for clarity, assume that *C* is a binary variable representing two possible states or outcomes (0 and 1), and consider the case *C* = 1 (these assumptions are relaxed later on). With this in mind, evaluate Equation 11 for *C* = 1, substitute the first term on the right side using Equation 12, and obtain

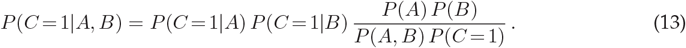

Next, note that the joint probability in the denominator of this expression can be written as

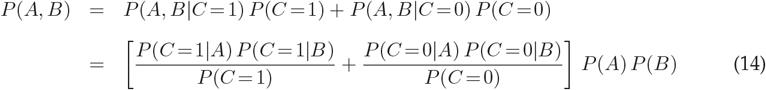

where Equation 12 was used twice in the last step, once for each of the conditional probabilities in the first step. Finally, combining Equations 13 and 14 gives

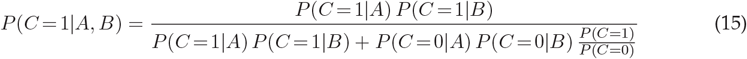

which is the fundamental result. It is stated for *C* = 1, but it is entirely analogous for *C* = 0 (the *C* = 1 and *C* = 0 terms simply exchange places). The more general way of writing this same result is this

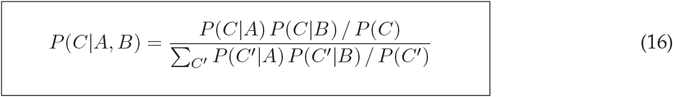

where the variable *C*^′^ is the same as *C*, except that it applies only to the sum in the denominator. These expressions provide a recipe for determining the probability of event *C* given two sources of evidence, *A* and *B*, and knowledge of how each of them predicts *C* separately, assuming that *A* and *B* act independently of each other with respect to *C*.

For the example above (Fig 1), the prior probability that Carol is home is *P* (*C* = 1) = 0.4. This is the estimate of her status without any further knowledge or measurement. The probability that Carol is home given that it is past 6 pm is *P* (*C* = 1|*A* = 1) = 0.6, and the probability that Carol is home given that the dog is not barking is *P* (*C* = 1|*B* = 0) = 0.6. Therefore, because the time and the barking are conditionally independent, it is straightforward to apply these numbers to Equation 15 to infer the probability that Carol is home given that, both, it is past 6 pm and the dog is not barking: *P* (*C* = 1|*A* = 1, *B* = 0) = 27*/*(27 + 8) ≈ 0.77. Certainty about her status grows when the two sources of evidence are in agreement, as one would expect, and Equation 15 tells exactly by how much.

In this example, the full frequency tree (Fig. 1) was given, which is to say that the three-way joint probabilities *P* (*A, B, C*) were all known. But a critical aspect of the above result is that it is applicable when this is not the case; it provides a way to estimate the outcome variable *C* when only the two-way relationships *P* (*C*|*A*) and *P* (*C*|*B*) are known. To understand the possible applications of Equation 15 (or 16), it is useful to briefly discuss its analytical properties.

### Key properties of the estimation formula

One way to view the implications of conditional independence is to consider the odds of the two outcomes for event *C*. In general, the odds given knowledge of *A* and *B* depend on two three-way measurements,

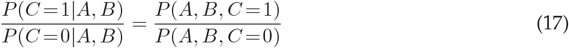

which are the two joint probabilities on the right. The standard way to deal with this calculation is to consider two ratios, one involving the likelihood of the observed data (the values of *A* and *B*) given the possible outcomes, and another depending on the priors for the outcomes

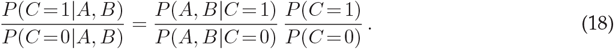

From here, the computational advantage afforded by conditional independence is easy to appreciate: the first term on the right can be factorized by direct application of Equation 6, such that

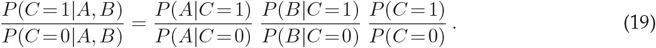

This step simplifies things by decoupling *A* and *B*. However, note that the outcome variable on the left, *C*, appears as a condition on the right. Under some circumstances, it might be easier or more intuitive to consider an equivalent factorization where *C* is always the outcome variable. That is,

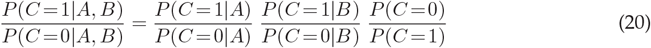

which results by application of Bayes’ rule to all the conditional probabilities on the right side of Equation 19. This expression is a natural counterpart to Equations 15, 16 (as can be seen by using either equation to evaluate the odds ratio on the left side). It gives the same result as Equation 19, but does so by parsing the effect of *A* on *C*, the effect of *B* on *C*, and the prior of *C* — the latter in inverted form compared to Equations 18 and 19.

Seen this way, it is clear that the events *A* and *B* are uninformative when they predict *C* with the same odds as the prior. When one of them is informative and the other is not (e.g., if *P* (*C* = 1|*B*) = *P* (*C* = 1) and *P* (*C* = 0|*B*) = *P* (*C* = 0) in Equation 20), two of the three factors cancel out and the odds for *C* simply adhere to those given by the single informative event. Because changes in outcome probability due to knowledge of *A* and *B* are relative to the prior, what matters most in this formulation are the *deviations* from the prior.

This gives rise to an interesting insight: for small deviations from the prior, the arithmetic for combining conditional probabilities is simply linear. This can be seen by setting

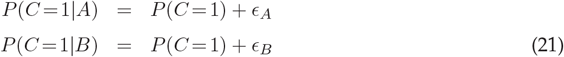

where *P* (*C* = 1) is sufficiently far from 0 and 1, and *ϵ*_*A*_ and *ϵ*_*B*_ are small positive or negative numbers with magnitudes ≪ 1, which means that events *A* and *B* add just a small amount of certainty. Then, according to Equation 15, the probability given both events is

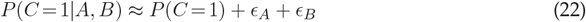

Thus, knowledge of *A* and *B* may shift the probability of a given outcome in the same direction (if the two deviations have the same sign) or may cancel each other (if the two deviations have different signs). In this regime, the contributions from *A* and *B* are simply added or subtracted.

In the earlier example (Fig. 1), the prior probability for Carol being home was *P* (*C* = 1) = 0.4; the conditional probabilities given each of two events separately, *P* (*C* = 1|*A* = 1) and *P* (*C* = 1|*B* = 0), were both equal to 0.6; and the predicted probability when both events were known was equal to 0.77 based on Equation 15. In turn, considering that the two corresponding deviations from the prior are equal to 0.2, the predicted probability is equal to 0.8 based on the linear approximation (Equation 22). The discrepancy is modest (*<* 5%) even though the deviations are substantial.

There is another aspect of Equation 15, also related to the unique arrangement of the prior terms, that may seem paradoxical. It would appear that, if *P* (*C* = 1) approaches 0, the second term in the denominator vanishes and the result for *P* (*C* = 1|*A, B*) must necessarily approach 1. However, this does not actually happen, because the remaining terms, *P* (*C* = 1|*A*) and *P* (*C* = 1|*B*), would also tend to 0; the prior and the two conditional terms are not independent of each other, so if the prior is small, the other terms must be as small or smaller. To see this, first note that the prior must be numerically equal to sums of joint probabilities involving either *A* or *B*, such that

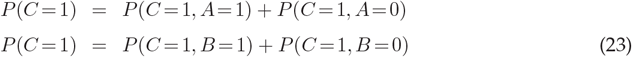

are both true. Then rewrite Equation 15 by multiplying the numerator and denominator by *P* (*A*)*P* (*B*)*/P* (*C* = 1). The resulting expression,

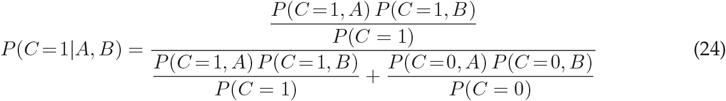

shows that the ratio in the numerator can never diverge to infinity: because of Equation 23, and because probabilities are ≥ 0, the joint probabilities *P* (*C* = 1, *A* = 1) and *P* (*C* = 1, *B* = 1) must be smaller than *P* (*C* = 1), or at most equal, so the ratio cannot exceed 1. By the same argument, the ratio involving *P* (*C* = 0) cannot diverge either. The paradox is resolved.

Finally, Equation 16 can be used to predict the state of the variable *C*, but one can also think of it as an alternative, operationally distinct way to define conditional independence. To investigate the conditions under which predictability implies conditional independence, start by assuming that Equation 16 is valid. Then apply Bayes’ rule to all the conditional probabilities in it so that each one is conditioned on *C*. After rearranging terms, the result is

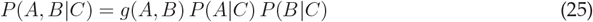

where the first term on the right hand side is defined as

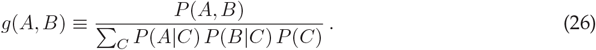

Equation 25, like Equation 16, is valid regardless of the number of states that the event variables can take, including in particular the number of states for *C*. And importantly, the case *g*(*A, B*) = 1 makes Equations 25 and 6 identical. Therefore, Equation 16 implies conditional independence when Equation 25 is valid and it follows that *g*(*A, B*) = 1. (All this excludes the case in which either *P* (*A*|*C*) = 0 or *P* (*B*|*C*) = 0, which is trivial) As it turns out, predictability (Equation 16) implies conditional independence under fairly general conditions — but not always. The equivalence depends on the dimensionality of the variables *A, B*, and *C*. If the number of values, or states, for these variables are *N*_*A*_, *N*_*B*_, and *N*_*C*_, then the critical condition is

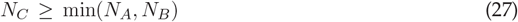

where min(*x, y*) is the minimum of *x* or *y*. When this condition is satisfied, Equation 16 implies conditional independence because the only way for Equation 25 to be true is if *g*(*A, B*) = 1. This can be seen as follows. First, multiply both sides of Equation 25 by *P* (*C*) so that

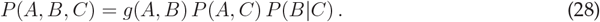

Then sum over all values (or states) of *B* on both sides,

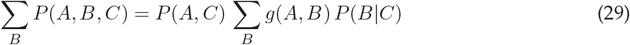

and notice that the sum on the left side is equal to *P* (*A, C*), which can be eliminated from both sides to yield

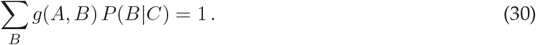

Similarly, using the same procedure but summing over all the values of *A*, it also follows that

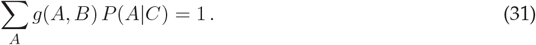

The next step is to rewrite the last two expressions using matrix notation, where indices denote the different states of the event variables *A, B*, and *C*. That is,

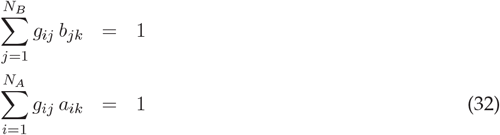

where *b*_*jk*_ is the probability *P* (*B* = *B*_*j*_|*C* = *C*_*k*_), which is the probability that *B* is in its j’th state given that *C* is in its k’th state, and so on for the other arrays. Using matrices, this pair of equations can also be written as

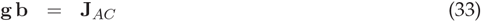

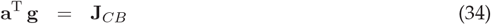

where J_*AC*_ is a *N*_*A*_ × *N*_*C*_ array of ones, J_*CB*_ is a *N*_*C*_ × *N*_*B*_ array of ones, and a^T^ is the transpose of a. We know that these two equations are satisfied when all the elements of g are equal to 1, because the probabilities *P* (*A*|*C*) and *P* (*B*|*C*) must add up 1 when summed over their argument; in other words, ∑_*i*_ *a*_*ik*_ = 1 and ∑_*j*_ *b*_*jk*_ = 1. The question, then, is whether the solution is unique. If *g*_*ij*_ = 1 is the only solution, then Equation 16 implies conditional independence (via Equation 25). Otherwise, Equation 16 represents a weaker constraint.

In general, Equations 33, 34 have a unique solution, *g*_*ij*_ = 1, if the inequality in Equation 27 is true; that is, if the number of states of the outcome variable *C* is at least as large as the number of states of *A* or *B*. This is why. In the arrays J_*AC*_ and J_*CB*_, all the rows and columns are the same. Therefore, to understand the possible solutions of the system, it suffices to consider a single row (for Equation 33) or a single column (for Equation 34) of the matrix g. For, say, the latter case, any solution must satisfy

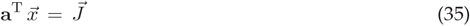

where 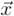 is an *N*_*A*_×1 column vector from g and 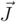 is an *N*_*C*_×1 column vector from J_*CB*_, with ones only. This is a classic linear system of equations where the elements of 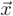 are unknowns. In such a system, a unique solution is typically possible if the number of equations, *N*_*C*_ in this case, is larger than or equal to the number of unknowns, *N*_*A*_ in this case. Otherwise, when the number of unknowns is larger, the system is underdetermined and an infinite number of solutions is possible. If the solution is unique, it must be the one we already know of, 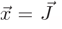, and it must be the same for all the columns of g; otherwise, if the solution is not unique, the columns of g can be different. Technically, the condition for a unique solution depends on the rank of the *N*_*C*_ ×*N*_*A*_ matrix a^T^, but this should be equal to min(*N*_*C*_, *N*_*A*_) unless a^T^ has linearly dependent rows or columns, which would correspond to redundant or trivial states for *A* or *C*. So, unless the probabilities in a^T^ have such an anomaly, the solution to Equation 34 is unique when *N*_*C*_ ≥ *N*_*A*_. Furthermore, by exactly the same argument, the solution to Equation 33 is unique when *N*_*C*_ ≥ *N*_*B*_, and because the solution g must be the same for both equations, either condition is sufficient. This is why the condition in Equation 27 is relative to the smallest between *N*_*A*_ and *N*_*B*_.

Therefore, if Equation 25 is true and the inequality in Equation 27 is satisfied, it must be the case that *g*(*A, B*) is equal to 1 for all combinations of *A* and *B*, which implies that these two variables are conditionally independent given *C*.

In summary, under fairly general conditions (namely, Equation 27), Equation 16 is valid if and only if *A* and *B* are conditionally independent with respect to *C*. The applications considered later on illustrate the use of Equation 16 (and extensions derived below) in both directions — as a recipe for generating predictions based on limited data, and as a method for characterizing the statistical associations between three variables.

### Conditional independence does not imply causality

The concept of conditional independence plays a central role in so called probabilistic graphical models [4, 9, 10], which consist of networks where each node represents a random variable and a connection between two nodes represents a statistical association between them. These statistical models are popular in part because they are a convenient platform for representing causal relationships between variables. It is important to realize, however, that the link between causality and conditional independence is not intrinsic. Just as correlation does not imply causation, in the case of three variables conditional independence does not imply any particular causal structure either; the analogy is appropriate, as there is a close relationship between conditional independence and partial correlation [11].

We illustrate this point by contrasting two examples with different causal structures. In the context of probabilistic graphical models (directed graphs, specifically), the two cases would be considered qualitatively different [4, 9, 10]. However, for the purpose of estimating the outcome of event *C* (via Equation 16), the distinction is immaterial.

The first case is when event *A* causes event *C* and event *C*, in turn, causes event *B*. This is the situation depicted in Fig 1, which we have discussed already: Carol is home (event *C*) depending on the time of day (event *A*), and the dog barks (event *B*) depending on Carol’s presence. The time of day and the dog barking are conditionally independent with respect to Carol’s presence, so the latter can be predicted based on limited information, namely *P* (*C*|*A*), *P* (*C*|*B*), and the prior *P* (*C*). Notably, conditional independence is satisfied simply because the numbers of observations across branches follow certain proportions.

Now consider the second case, which is as follows. Suppose that event *C* indicates whether in the town of Wreckville a car crashes (*C* = 1) or does not crash (*C* = 0) within a year. The pertinent conditions are whether the car is antiquated (*A* = 1) or not (*A* = 0), and whether the driver has a blameless record (*B* = 1) or not (*B* = 0). Say that the the overall probability of a crash, the probability of a crash given that the car is antiquated, and the probability of a crash given that the driver does not have a blameless record are

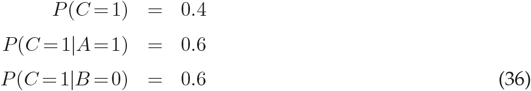

and the question is, what is *P* (*C* = 1|*A* = 1, *B* = 0), i.e., what is the probability that a car crashes within a year given that it is antiquated and that the driver’s record is not blameless? In this case, events *A* and *B* are both causal to event *C*, but the exact answer is still given by Equation 15 provided that *A* and *B* are conditionally independent with respect to *C*. Because both the poor state of the car and the track record of the driver increase the risk of an accident, the sought probability should be larger than 0.6; and indeed, Equation 15 gives 27/35 ≈ 0.77 in this case.

If these numbers seem familiar, it is because they are exactly the same as in the case of Carol’s barking dog. In fact, the full frequency tree in Fig 1 is applicable to this example; the only thing that changed were the verbal labels attached to the variables *A, B*, and *C*, and their corresponding mental associations. Given a data set and the task of estimating the outcome of a variable, what matters is whether conditional independence is numerically satisfied; the particular causal structure ascribed to the variables neither guarantees nor negates this.

The different causal interpretations of the same data set also reinforce an observation made earlier, that conditional independence (between *A* and *B* with respect to *C*) and plain independence (between *A* and *B*) are orthogonal properties. In the case of the car crash, it is tempting to think that a crash is predictable because the state of the car and the experience of the driver are independent of each other, but this would be wrong; as remarked before, variables *A* and *B* deviate substantially from independence given the numbers in Fig 1. Rather, their key virtue is that they act independently with respect to *C*. This is what conditional independence establishes.

The traditional view of conditional independence describes how the variable *C* affects the association between *A* and *B*; specifically, when the condition is valid, their joint occurrence conditioned on *C* can be factorized (right side of Equation 6). In contrast, the new results take the reverse view; although the factorization is more complicated (right side of Equation 16), conditional independence now tells us something about how *A* and *B* affect *C*, in a statistical sense.

### Expected accuracy in higher dimensions

Equation 16 is valid for event variables *A, B*, and *C* that can take any number of values, or states, not just two. An important extension, however, is to the case where more than two sources of evidence may be predictive of event *C*. For example, in addition to the time of day and the dog barking, the presence of a car in the garage may also be informative of whether Carol is home or not.

First consider the binary case, in which *C* is either 0 or 1 (Equation 15). If instead of *A* and *B* there are *N* events *E*^(1)^, *E*^(2)^, …, *E*^(*N*)^, all of which are conditionally independent given *C*, then the probability that *C* is in state 1 given knowledge of all those events is

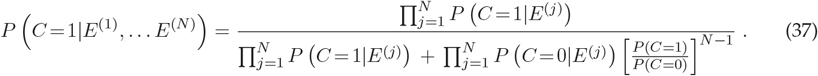

With *N* = 2, this equation is the same as Equation 15. It is also written for one particular outcome, *C* = 1, but the probability of the alternative outcome is entirely analogous.

The most general expression allows for both multiple sources of evidence, as above, and multiple states of the variable of interest, as in Equation 16. If *C* can take any of *M* possible values *C*_1_, *C*_2_, …, *C*_*M*_ and the observed events *E*^(1)^, *E*^(2)^, …, *E*^(*N*)^ are conditionally independent across all of them, the probability of a particular outcome *C*_*j*_ given the *N* observed events is

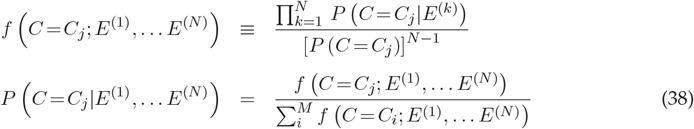

where the denominator in the last equation is simply a normalization factor. This is the probability that the variable of interest is in state *j* (one of *M* possible states) given that *N* conditionally independent sources of evidence are known. This is the main analytic result. Equations 16 and 37 are lower-dimension versions of this expression (with terms arranged slightly differently). In this form, it is clear that what matters is how each conditional probability *P C* = (*C*_*j*_|*E*^(*k*)^) deviates from the prior, *P* (*C* = *C*_*j*_), in terms of their ratio.

The main point is that these equations provide a statistical expectation for what should happen if multiple sources of evidence act independently given the quantity of interest, *C*. This provides a parameter-free benchmark for analyzing the efficacy of evidence-integration processes in a variety of situations, as discussed in the rest of the paper.

### Repeated diagnostic tests for a disease

A paradigmatic situation in which probabilities are inverted, as in Equation 11, is in clinical settings, when a diagnostic or screening test is applied to detect a disease. A central question is how to combine the results of multiple diagnostic tests, and in this context it has long been recognized that conditional independence may be critical because it greatly simplifies all the calculations involved, especially when there is no gold standard for determining the true disease state [12, 13, 14]. Here we consider a scenario in which a gold standard does exist; in that case, it is possible to base all the calculations directly on the predictive values (defined below) of the individual tests, which are the most relevant quantities for diagnosis. This example is useful because it considers the derived results in a familiar context, and clearly illustrates how they can be used to (1) predict an outcome of interest based on limited information, or (2) test whether conditional independence is satisfied, which is necessary for establishing whether making such predictions is warranted. The two applications are discussed in that order.

Let the variable *D* represent the presence (*D* = 1) or absence (*D* = 0) of a disease, and let the variable *T* represent the outcome of the test, which can be positive (*T* =+) or negative (*T* =−). In this context, two probabilities characterize the effectiveness of the test,

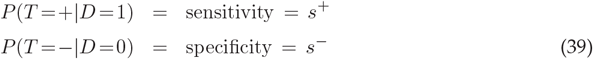

where *s*^+^ and *s*^−^ are just abbreviations for these quantities, which are known as sensitivity and specificity [15, 16]. Knowing these two probabilities, together with the prevalence of the disease, which is its prior probability, *P* (*D* = 1), allows one to use the test as a diagnostic; that is, it makes it possible to test a new patient and determine the probability that the disease is indeed present when the test is positive,

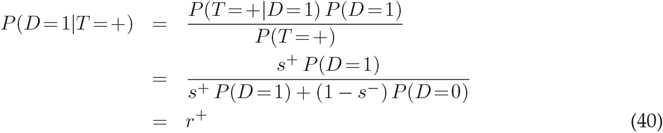

or the probability that the disease is indeed absent when the test is negative,

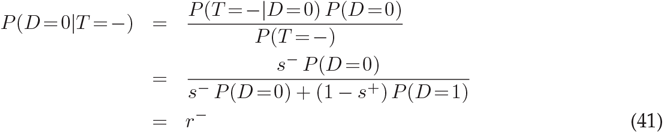

where *P* (*D* = 0) is simply 1−*P* (*D* = 1) and, again, *r*^+^ and *r*^−^ are simply abbreviations for the probabilities defined on the left-hand sides of these equations. These probabilities are often referred to as the positive predictive value (*r*^+^) and the negative predictive value (*r*^−^) of the diagnostic test [17].

Now suppose that the test can be repeated multiple times, for instance by drawing multiple samples from the same patient [18]. In that case, if the outcomes of these repeated tests are conditionally independent given the disease state, *D*, then Equation 37 can be used to determine the probability that the disease is present given that *n* out of *N* samples turn out to be positive. Because all the test repetitions have the same sensitivity and specificity, and thus the same posterior probabilities *r*^+^ and *r*^−^, Equation 37 becomes

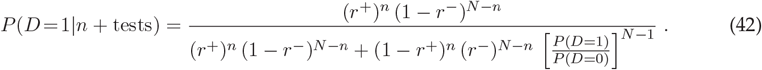

The behavior of this expression for different values of *n* and *N* demonstrates the power of combining conditionally independent evidence samples (Fig 2). Suppose that both *r*^+^ and *r*^−^ are relatively close to 0.5, so the test is rather unreliable and a single application provides little information (Fig 2a, top). If the test is applied 4 times (Fig 2a, second row), its accuracy increases substantially when all 4 tests are positive or all are negative, but the procedure is still not very useful because for the most common outcomes (1 or 2 positive tests; see dashed lines) the resulting probabilities still do not deviate from 0.5 very much. However, as the test is repeated more times, the results eventually become both accurate (probabilities close to 1 or 0) and unambiguous, as the range in which the number of positive tests provides an uncertain diagnosis becomes progressively less likely (Fig 2a, bottom plot; note little overlap between dashed lines).

**Fig 2.**
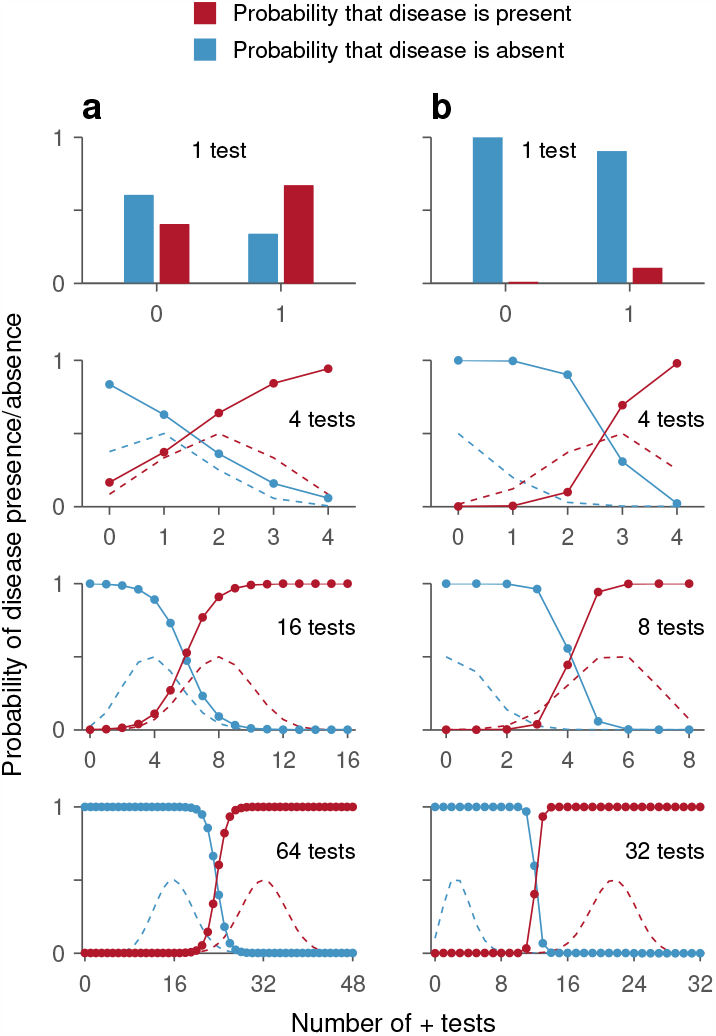
Multiple conditionally-independent repetitions of a diagnostic test increase diagnostic accuracy. The plots show results from Equation 42 for different repetitions of a test (*N*, varying across rows) as a function of the number of positive tests obtained (*n*, along the x-axes). Red bars and data points correspond to *P* (*D* = 1|*n* + tests); blue bars and data points correspond to *P* (*D* = 0|*n* + tests). Dashed lines are proportional to the likelihood of obtaining a particular number of positive tests when the disease is present (red) or absent (blue). **a**, Example where a single test has low sensitivity and specificity. In this case, *s*^+^ = 0.5, *s*^−^ = 0.75, and *P* (*D* = 1) = 0.5, so one test is relatively uninformative. **b**, Example where a single test has low sensitivity and high specificity. In this case, *s*^+^ = 0.67, *s*^−^ = 0.91, and *P* (*D* = 1) = 0.0148, so a single negative test is clearly informative but a positive one not so much. With increasing repetitions, the diagnosis becomes more accurate and less ambiguous.

Such convergence to high accuracy is also observed when one test result (positive or negative) is much more informative than the other. In an example that approximates the case of a rare disease, i.e., one with relatively low incidence, a single negative test indicates that the disease is very likely absent, but a single positive test provides negligible evidence that the disease is present (Fig 2b, top). However, in spite of the strong asymmetry, multiple repetitions of the test eventually yield accurate and unambigous diagnoses for both disease presence and absence (Fig 2b, bottom) — as long as the tests remain conditionally independent of each other given the disease state.

This example illustrates how to generate a prediction for the performance of a detector based on multiple repetitions of a test when the conditional independence between replicates of this test is either assumed or known to be applicable. However, the same relationship (Equation 42) can be used to determine whether multiple repetitions of a test are, in fact, conditionally independent given the state of the tested individual, or to quantify the degree to which the data deviate from conditional independence.

Suppose that the repeated-test protocol is applied many times and that the ground truth about the presence or absence of the disease can be determined by a separate gold-standard test. In that case, the predictive values (i.e., posterior probabilities) can be determined empirically from the observed results. For instance, for a four-test protocol one would compute the probability that the disease was actually present given that 0, 1, 2, 3, or 4 tests were positive. These measured probabilities would then be compared against those predicted based on the single-test data and the assumption of conditional independence, via Equation 42 (i.e., the data in Fig 2, row 2). For repeated tests that are indeed conditionally independent given the disease state, a plot of the empirical probabilities against the predicted ones would produce data points lying along the main diagonal. In contrast, for repeated tests that violate the independence condition, the data points would deviate from the diagonal, and the magnitude of the deviations would be indicative of the degree of correlation between the repeated tests (Fig 3).

**Fig 3.**
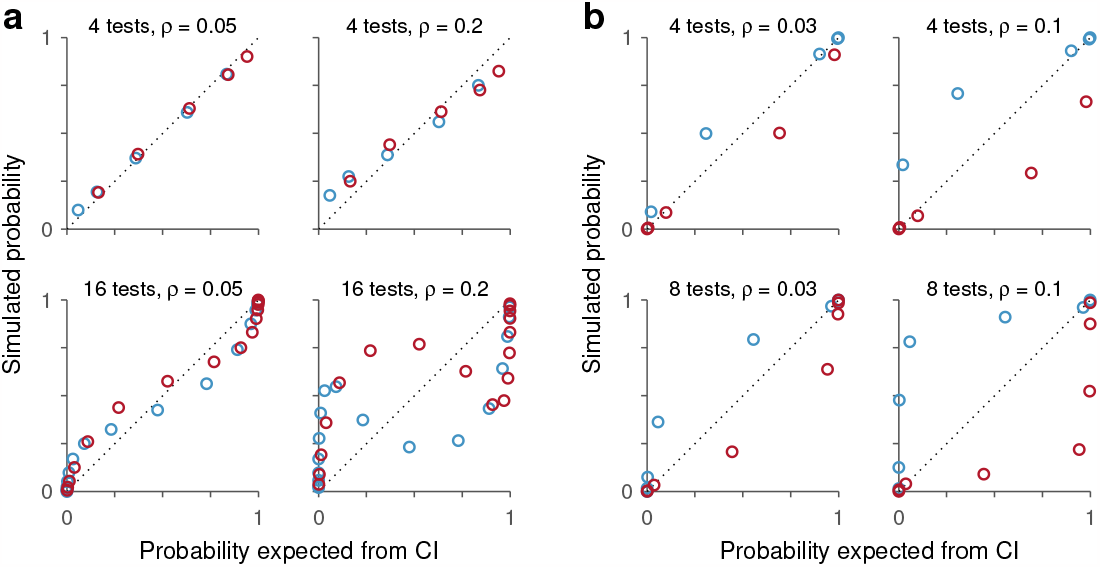
Determining conditional independence across repetitions of a diagnostic test. The plots compare the results obtained from Equation 42 (x axes), which are based on test outcomes that are conditionally independent given the disease state, with the results of simulations (y axes) in which the test outcomes were correlated. The number of repeated tests is indicated in each case, along with their correlation coefficient (*ρ*). Points correspond to *P* (*D* = 1|*n* + tests) (orange) and *P* (*D* = 0|*n* + tests) (blue). **a**, Example where a single test has low sensitivity and specificity (same as in Fig 2a). **b**, Example where a single test has low sensitivity and high specificity (same as in Fig 2b). The discrepancy between the independent and non-independent conditions increases with the number of test repetitions and their degree of correlation (*ρ*). CI, conditional independence.

To illustrate the possible results, a subset of the conditions depicted in Fig 2 were simulated with diagnostic tests that, upon repetition, were correlated to varying degrees (Fig 3; Methods). We found that the discrepancy between the predicted and “empirical” results can vary widely depending on the specific conditions of the simulation. Of course, a higher degree of correlation produced differences of larger magnitude (Fig 3, compare left versus right columns in panels a and b), as one would expect. However, the impact of the correlation was milder for fewer repetitions than for more (Fig 3a, b, compare top versus bottom rows), and depended on the symmetry of the sensitivity and specificity values as well (Fig 3, compare a versus b). We also found that the results depended only weakly on the exact method used to generate the correlations (Methods), which in real data may arise for many reasons. The examples suggest that, under certain conditions, the assumption of conditional independence may be a reasonable approximation even when it is not strictly true.

### Calculating biological age

There is much interest in understanding the fundamental processes that lead to performance decline during aging; for instance, how environmental versus genetic factors contribute to it. An important concept in this context is that of biological age, which represents the functional state of a biological system along its lifelong trajectory irrespective of its chronological age [19]. Practical determination of biological age is thought to be potentially useful as a more accurate predictor of death or disease than standard chronological age [20].

Numerous physiological processes have been identified that evolve in stereotypical ways over the normal course of aging. Some of these aging biomarkers are widespread molecular processes [21, 22]. For instance, telomere length is indicative of the number of cell division cycles that the organism has gone through; DNA methylation is indicative of gene transcription activity and serves as an “epigenetic clock”; the levels of certain proteins are indicative of cellular senescence and correlate strongly with chronological age; etc. Many other physiological markers of aging are organ-specific [19]. For example, lung function as measured by peak aerobic capacity is one of the most reliable correlates of aging, but cardiovascular, immune, neurocognitive, and other systems show characteristic trajectories with aging that have both macroscopic and cellular manifestations. Knowing the actual capabilities of these systems at a particular moment, i.e., their effective wear and tear, so to speak, can serve to define the biological age of a subject. For our purposes, the key question is how. How should multiple aging biomarkers be combined into one quantity that corresponds to biological age?

There are several possible approaches [23, 24, 25, 26], but Equation 38 provides a path that is potentially data efficient and leads to a simple, easily interpretable result. Furthermore, because many of the identified biomarkers of aging relate to widely different biological processes, at least a subset of them are likely to be conditionally independent with respect to chronological age. We illustrate this with a hypothetical example (Fig 4).

**Fig 4.**
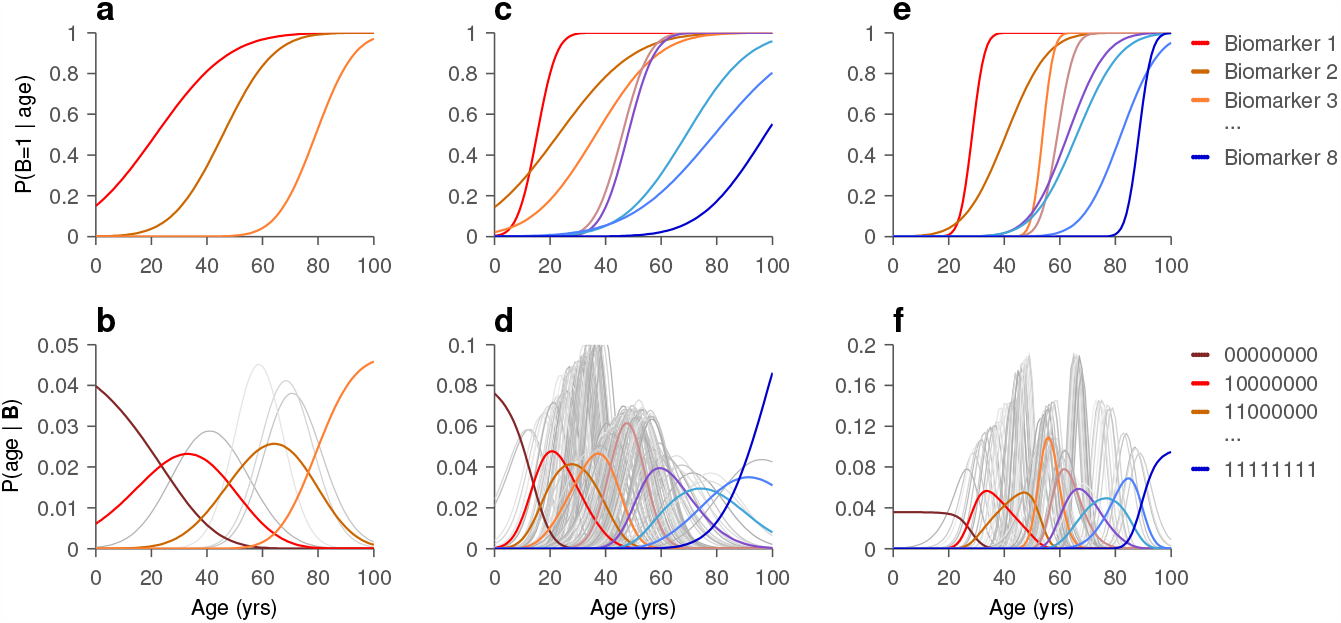
Estimation of biological age. **a**, Probability curves for three hypothetical biomarkers of age. Each biomarker is assumed to have two possible states, and each plotted function corresponds to *P* (*B* = 1|age). **b**, Each curve shows the probability that a subject has a particular age given the measured states of the three biomarkers above, assuming that they are conditionally independent of each other with respect to age. The 8 curves plotted correspond to the 8 possible states of the 3 (binary) biomarkers. The left-most curve (dark red) corresponds to *P* (age|000), which is when all biomarkers are in the 0 state; the next curve (bright red) corresponds to *P* (age|100), which is when the first biomarker is in state 1 and the other two are in state 0, and so on. Colored curves are for congruent states, in which the biomarkers are activated in an order consistent with the midpoints of their probability curves (states 000, 100, 110, and 111); light gray curves are for the remaining, incongruent states (states 010, 001, 011, 101). **c, d**, As in **a, b**, but with 8 biomarkers. Now 256 curves are plotted in panel **d**, 8 congruent and 248 incongruent. **e, f**, As in **c, d**, but with biomarkers that generally switch state more abruptly as functions of age.

Consider three aging biomarkers, each of which can be in two states, 0 or 1, and suppose that the probability of being in state 1 increases with age (Fig 4a, top). One such probability function could describe, for instance, the fraction within a given subject population for which telomere length has dipped below a certain threshold, or the fraction for which the concentration of a particular hormone is below a given cutoff, etc. Whether the functions increase or decrease does not matter; they do not even need to be monotonic. In the examples, all the biomarker functions are shown as monotonically increasing simply for ease of visualization, but what is important is that they represent probabilities. With this in mind, the expression

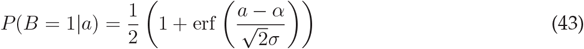

where erf is the error function, corresponds to the probability of biomarker *B* being in state 1 at each value of *a*, which represents chronological age, and this probability is characterized by two biomarker-specific parameters: *α*, which determines the midpoint of the curve, and *σ*, which determines the steepness of its rise. In the examples below with three or eight biomarkers, each one follows an equation like this but with different parameter values, which were drawn from distributions chosen to produce a diversity of biomarker response curves (Fig 4a, top).

Another necessary ingredient for the computation of biological age is knowledge of the overall prior age distribution, *P* (*a*). For illustration purposes, we consider the age distribution to be flat between 0 and 100 years, with zero probability elsewhere. This simplifies the discussion, but is by no means a requirement and does not affect the conclusions.

Having defined the prior age distribution, *P* (*a*), the prior for each biomarker state, i.e., *P* (*B* = 1) and *P* (*B* = 0), can be obtained by combining *P* (*a*) with Equation 43. Then, Equation 43 can be inverted according to Bayes’ theorem to generate the probability function *P* (*a*|*B*) for each biomarker. Finally, using Equation 38, we can compute the probability that a subject has a particular age given knowledge of the three biomarkers. With 3 binary biomarkers there are a total of 2^3^ = 8 possible state vectors, so 8 possible curves describing the likely values of age given the measured physiological state (Fig 4b). For instance, when the state vector is 000, none of the biomarkers have transitioned to state 1, so the subject is likely to be very young (Fig 4b, dark red, leftmost curve). When the state vector is 110, the first two biomarkers have transitioned but the third has not, so the subject is most likely to be about 65 years old (Fig 4b, gold curve, third colored curve from left). Some state vectors are more likely (Fig 4b, colored curves) than others (Fig 4b, gray curves), but they all lead to a distribution of ages that reflects the measured condition of the subject. By this procedure, biological age is defined as the age that would be inferred given the physiological condition of the subject in reference to comparable historical data.

The situation is entirely analogous when, say, 8 biomarkers instead of 3 are used (Fig 4c, e), except that now 2^8^ = 256 state vectors are potentially observable (Fig 4d, f, note the number of curves). With more biomarkers, the distributions of likely ages given the states are typically narrower than with fewer biomarkers (Fig 4, compare b versus d), but the spread of these resulting distributions also depends on the steepness of the biomarker state transitions (Fig 4, compare c versus e, and d versus f).

The point of this example is to show that, having identified a set of physiological quantities that vary reliably with age, and having formulated their dependencies in terms of state-transition probabilities, there is a natural way to define biological age — namely, as the age that one would infer based on historical data, a subject’s current physiological condition, and the rules of probability. And this inference is particularly easy to compute when the physiological quantities are conditionally independent of each other with respect to age.

As with the example in the previous section, the same equations can be used to infer a quantity of interest (age in this case) or, alternatively, to test the degree to which the measured variables (the biomarkers) deviate from conditional independence. To illustrate the latter process, we simulated the measurement of the 8 binary biomarkers with probability curves shown in Fig 4c. For a given age and a given iteration, the 8 responses were generated by comparing 8 randomly drawn samples from a uniform distribution to the corresponding probability-curve values, as is customary for simulating probabilistic outcomes. However, these random samples were correlated (Methods). As a result, the biomarker readouts for each iteration — that is, for each simulated subject — were also correlated. This way, it was possible to produce non-independent biomarker readings, as shown by the resulting matrix of correlation coefficients (Fig 5a); when one biomarker was in state 1, other biomarkers from the same simulated subject were more likely to be in state 1 too. That the resulting biomarkers were not conditionally independent with respect to age can be verified by comparing the probability of any given biomarker state vector B at each age in two cases: as expected from the probability curves and conditional independence (Fig 5b, x axis), and as observed from the actual simulated biomarker states (Fig 5b, y axis). Although the data points do not fall too far off the diagonal, the underlying correlations generated state-vector probabilities that clearly deviate from those obtained with zero correlation. More importantly, however, the correlations only caused a relatively modest distortion of the posterior probability for age given the state vector. That is, the age distributions inferred given the actual, correlated biomarker state vectors (Fig 5c, y axis) were fairly similar to the distributions obtained by ignoring those correlations and assuming conditional independence (Fig 5c, x axis). The results were qualitatively the same for different numbers of biomarkers (from 3 to 12) and regardless of the specific parameter values used (*α* and *σ* in Equation 43).

**Fig 5.**
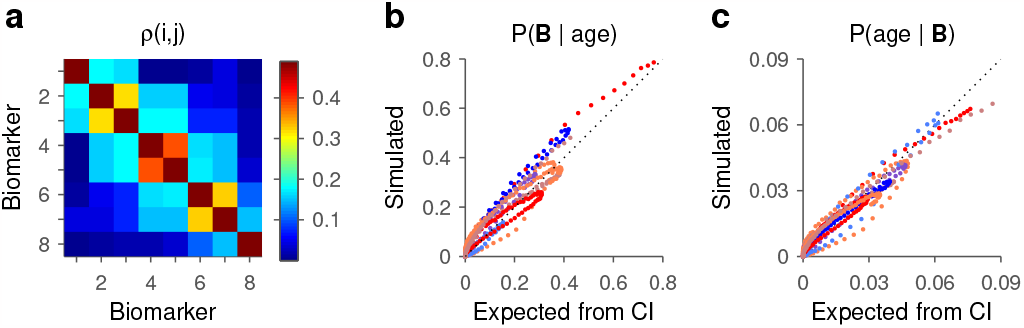
Determining conditional independence between biomarkers of aging. Results are for the same set of 8 binary biomarkers depicted in Fig 4c, but either satisfying or violating the conditional independence assumption. **a**, Matrix of correlation coefficients obtained after simulating the correlated responses of the 8 biomarkers. For each biomarker pair, correlation values were computed at each age and results were then averaged across ages. Thus, the expected correlation would be zero under conditional independence. **b**, Data points indicate the probability of a biomarker state B given the age of the subject, either based on the conditional independence assumption (x axis) or on simulated biomarker outcomes (y axes) that were correlated as shown in **a**. Each curve of a given color corresponds to one biomarker state, and each point of that color corresponds to one age. Results are for the nine congruent state vectors, that is, vectors 00000000, 10000000, 110000000,… and 11111111. Same color scheme as in Fig 4d. Note deviations from the diagonal demonstrating the impact of the correlations between outcomes. **c**, Data points indicate the probability of a particular age given the biomarker vector state. Values on the x axis are as expected from conditional independence (Equation 38; same values as on the y axis of Fig 4d). Values on the y axis are as determined from the simulated biomarker states with correlations. With binary biomarkers, conditional independence provides a good approximation to the true posterior probability *P* (age|B) even when the assumption is not strictly true.

This example illustrates both applications of the expressions derived at the beginning in the context of an interesting biomedical problem, but still within a highly simplified scenario. Addressing the problem in reality would require much more detail, but this is a proof of concept. The point is that conditional independence is a potentially useful tool either for selecting informative biomarkers that are non-redundant, or as an alternative method for estimating biological age.

### Integration of sensory cues from multiple modalities

Everyday behaviors are often guided by diverse types of sensory input. For instance, hearing the muffled sound of broken leaves while gleaning a grayish object suddenly moving in the periphery may lead one to conclude that a squirrel just hid behind a tree. There is much interest in understanding the mechanisms whereby neuronal circuits combine multiple sources of information, as in this example, to yield a signal that is more reliable and informative than its individual components. When the individual cues consist of sensory stimuli of different qualities or modalities, this problem often falls under the rubric of multisensory integration [27, 28, 29].

In practice, cue integration studies often rely on behavioral tasks in which a choice must be made based on two cues — we will call them *A* and *V* in this case — and success in the task is measured for various combinations of cue values. A common design is to interleave trials in which *A* is presented alone, *V* is presented alone, and *A* and *V* are presented together (Fig 6a-c). A fundamental question in this context is whether the performance of the subject (or the response of a recorded neuron) in the multi-cue condition is higher than would be expected based on the performance levels seen in the single-cue conditions. Choice accuracy above such expectation may be indicative of active mechanisms dedicated to generating combined signals that are most effective or advantageous; and vice versa, performance below expectation may be diagnostic of internal correlations or other constraints that may limit the processing power of the underlying circuitry [27, 28, 29].

Interestingly, though, it is often unclear what the default expectation should be. Prior approaches have been at least in part tailored according to the measurement of interest, which may be choice accuracy (percent correct), reaction time, or the firing activity of a neuron, for instance. In behavioral studies, a popular benchmark is a Bayesian model that makes broad statistical assumptions about the underlying signal detection processes and relevant noise distributions; i.e., about the variability in the internal representations of the cues [30, 31, 32]. Alternatively, there have been various proposals for estimating the increase in performance that should be expected based simply on the fact that in the multicue condition the system has two sensory signals to ‘choose’ from, rather than one [33, 34, 35, 36, 37]. The idea in this case is that each sensory channel can independently drive the decision process, but at any point in time only the stronger signal contributes to the subject’s response. We submit that the performance expected when the two cues are conditionally independent relative to the choice provides a principled, parameter-free benchmark that in many cases is ideal for evaluating the effectiveness of multisensory integration. A recent study in mice [38] took a similar approach.

To illustrate this, we consider a simple hypothetical task that captures the essence of the problem. In the multisensory spatial discrimination task, a cue is given in each trial and the subject must indicate which side it originated from, left or right. Auditory (Fig 6a), visual (Fig 6b), and multisensory trials (Fig 6c) are randomly interleaved, and all stimuli are brief (say, 50 ms). The challenge is to detect the unisensory or multisensory cue, which can be very faint, but in the context of a two-alternative forced-choice task, for which the chance level is unambiguous. To demonstrate the full range of possible results, we assume that the luminance of the visual stimulus varies between 0 and a maximum value, such that performance in the unisensory visual trials covers the full range between chance (50% correct) and 100% correct (Fig 6d, black curve). The auditory stimulus is instead delivered at two intensities, both of them yielding performance levels in between the two extremes (Fig 6d, blue horizontal lines). The key question is: given the performance in the two unisensory conditions, what performance should be expected in the multisensory case?

Conditional independence provides a parameter-free answer that is consistent with the absence of a functional interaction between the two stimuli. When the visual and auditory signals are conditionally independent with respect to the subject’s choice, the expected performance curves (Fig 6d, light and dark red curves) demonstrate improved accuracy that stays within the limits imposed by the representation of probabilities. At the lowest luminance, multisensory choices are just as accurate as with the auditory stimulus alone; this makes sense because a visual stimulus that is entirely uninformative (thus yielding chance accuracy) cannot possibly improve performance when combined with an informative auditory cue. At the other end, when the visual stimulus is of the highest luminance, choice accuracy in visual-only trials is already near 100% correct, so an additional auditory cue cannot increase performance much further. Overall, in the multisensory conditions, the fraction of correct choices shows a graded increase with luminance that parallels that in the visual-only condition, but bracketed between the two limits just discussed.

The predicted multisensory curves in this example were generated from a formula that is essentially identical to Equation 15, but there is a subtle difference that is important to explain. Consider an auditory stimulus of fixed intensity and a visual stimulus of fixed luminance, and take *V* = 1 to mean that the visual cue was shown and *V* = 0 to mean that it was not. The quantity of interest is then *P* (*C* = 1|*A* = 1, *V* = 1), which is the probability of a correct outcome given that both cues were presented. Application of Equation 15 would seem straightforward, except that the probabilities on the right hand side of the equation do not match the probabilities obtained from the unisensory trials in the experiment. For instance, the term *P* (*C* = 1|*A* = 1) corresponds to the probability of a correct choice when we know that the auditory cue was shown and we *do not know* whether the visual cue was shown or not; in contrast, the unisensory auditory experiment yields *P* (*C* = 1|*A* = 1, *V* = 0), which is the probability of a correct choice when we know that the auditory cue was shown and that the visual cue was not. The same goes for the visual unisensory experiment. So, Equation 15 is not applicable to the unisensory data at hand.

However, the conditional probability for the multicue trials can be cast in terms of the probabilities obtained from the unisensory trials. To do this, first write down the four versions of Equation 20 (with *V* taking the place of *B*) that result when the four possible combinations of the *A* and *V* binary events are considered. Then combine those expressions to eliminate the terms conditioned on *A* alone or *V* alone. After some algebra, the result is

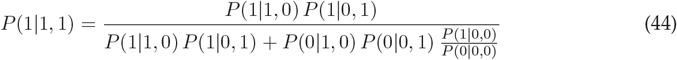

where we have used the abbreviations

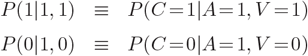

and so on, always assuming the same order, (*C*|*A, V*). Equation 44 has the exact same form as Equation 15 but with the terms on the right hand side correctly matching the knowledge obtained from the two unisensory experiments. Another notable aspect of this expression is that the terms that previously represented the priors for correct and incorrect choices, *P* (*C* = 1) and *P* (*C* = 0), have now turned into *P* (*C* = 1|*A* = 0, *V* = 0) and *P* (*C* = 0|*A* = 0, *V* = 0), which are the probabilities for correct and incorrect choices when neither cue is shown, which simply correspond to the chance levels of the experiment. The multisensory curves in this example (Fig 6d, light and dark red curves) were obtained by plugging the performance values in the simulated unisensory conditions into the above expression, along with 50% chance probabilities for correct and incorrect choices.

This case also serves as a reminder that, because conditional probabilities such as *P* (*C*|*A* = 1, *V* = 0) and *P* (*C*|*A* = 0, *V* = 1) represent empirical data and thus have some inherent uncertainty associated with their measurement, any estimate from Equation 44 (or versions discussed earlier) will also have some uncertainty. Importantly, with discrete random variables, as considered here, a distribution for the predicted value can be generated numerically by combining binomial statistics with knowledge of the numbers of observations involved in the measurements (Methods). An example is shown in the form of 68% confidence intervals around the multisensory predictions (Fig 6d, shaded ribbons), which were created by assuming that, for each of the two auditory experiments, the probability *P* (*C*|*A* = 1, *V* = 0) was based on 2000 observations or task trials, and for each luminance level in the visual experiment, the probability *P* (*C*|*A* = 0, *V* = 1) was based on 500 trials. Other statistics for the predicted probabilities (e.g., significance values) can be obtained from the same procedure.

Equation 44 is an alternative formulation of the conditional independence relationships derived earlier (Equation 15). Although the data that these two expressions use is slightly different, they give rise to the same bounded arithmetic between the combined probabilities. This latest version matches the typical structure of a multisensory experiment, wherein the performance observed with a multisensory cue is compared to that obtained with separate unisensory cues. This benchmark for multisensory interaction should be applicable under a wide range of conditions; namely, whenever (1) the primary measurements can be framed in terms of probabilities, and (2) chance performance is well defined within the experimental design.

The next step in a multisensory experiment like the one just discussed would be to compare the observed multisensory performance to the predictions from Equation 44. As mentioned above, empirical performance above the predictions would be indicative of synergistic interactions between the two cues that lead to a more robust internal signal, whereas empirical performance below the predictions would hint at the presence of deleterious correlations between the two sensory channels or other internal limitations to their optimal processing [39, 40, 41].

### How past events interact to influence ongoing performance

Much is known about the mechanisms that guide visuomotor behaviors. For instance, it is well established that the decision of where to look next depends in lawful ways on two factors, the salience of the objects in view, and the subject’s internal goals, e.g., what a person might be looking for, such as a pen on a desk, a face in a crowd, and so on [42, 43, 44]. However, more recently it has been recognized that there is a third, independent factor that also plays a key role in guiding our attention and eye movements, which is generally referred to as selection history [45, 46, 47]. This includes effects attributable to recent experiences, such as past stimuli seen or past rewards gained or lost, that do not directly affect goals and intentions. The signature manifestation of a relevant history variable is a difference in performance when the variable is repeated across trials (e.g., successive red targets) versus when it switches value (e.g., a red target after successive green targets). The final example illustrates the application of conditional independence to the problem of disentangling how different types of selection history interact to influence a subject’s behavior in a visuomotor task.

Consider a simple search task in which the objective is to locate the oddball [48, 49], the object that has a different color (Fig 7a). After the subject fixates a central spot, a cue stimulus is briefly displayed and then masked, and the subject is required to make an eye movement to the location where the oddball was presented. From trial to trial, the oddball position varies randomly between four possible locations (up, down, left, right) and its color varies randomly between red (with green distracters) and green (with red distracters). The cue presentation period is short enough, and the chromatic contrast between target and distracters small enough, that average performance is above chance (25% correct) but well below 100% correct.

**Fig 6.**
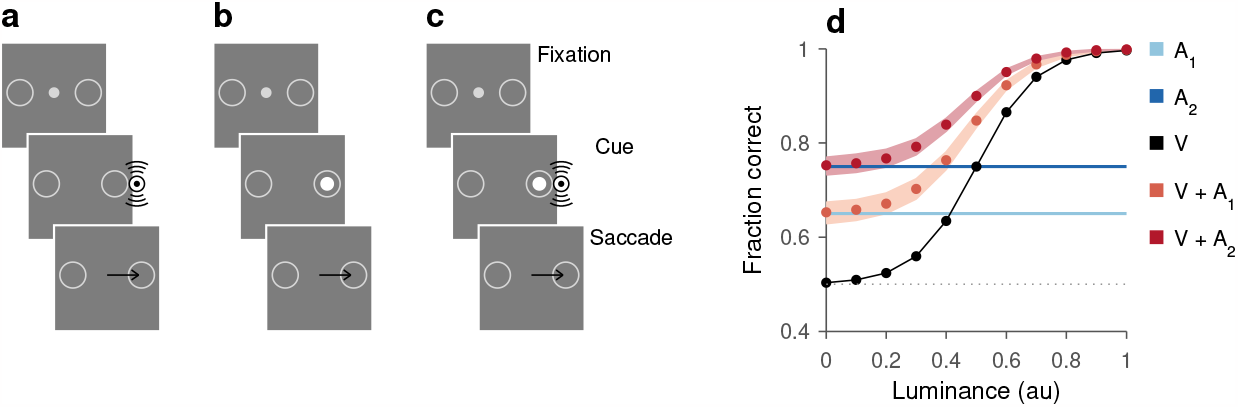
Conditional independence as a benchmark for performance in a hypothetical multisensory integration task. **a**–**c**, Sequence of events in the multisensory spatial discrimination task. After the subject fixates their gaze on a central point (Fixation), a cue stimulus is briefly presented (Cue) either to the left or the right of fixation. The subject’s task is to respond by making an eye movement (Saccade) to the choice target (open circle) that is on the same side as the stimulus. Three trial types are interleaved: unisensory auditory trials (**a**), unisensory visual trials (**b**), and multisensory trials (**c**). Black arrows indicate eye move-ments. **d**, Simulated fraction correct as a function of visual cue luminance in five experimental conditions. Blue horizontal lines mark performance in unisensory auditory trials with two sound intensities (A_1_ and A_2_). Black symbols indicate performance in unisensory visual trials (V). Light and dark red symbols indicate the multisensory performance expected from conditional independence when combining the visual cues with the first (V + A_1_) or the second (V + A_2_) auditory stimulus. Shaded ribbons indicate 68% confidence intervals for the predictions, obtained under the assumption that 2000 trials were run in each of the auditory experiments and 500 trials per luminance value were run in the visual (Methods). Chance performance is at 50% correct (dotted line).

**Fig 7.**
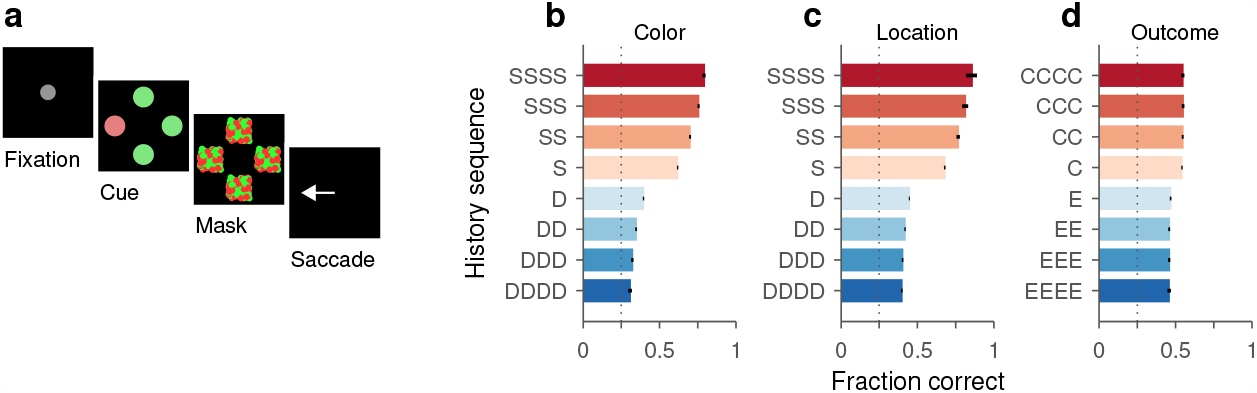
Selection history effects in a hypothetical oddball search task. **a**, Sequence of events in the oddball task. After the subject fixates their gaze on a central point (Fixation), the cue stimulus is briefly presented (Cue) and followed by a mask (Mask). The subject then indicates the location of the target with an eye movement (Saccade) and receives feedback indicating what the true location was (not shown). **b**, Fraction correct (x axis) for choices preceded by different color histories (y axis) going 1–4 trials back, with prior target stimuli having been of the same (S) or different (D) color as the target in the current trial. **c**, As in **b**, but for performance conditioned on the history of preceding target locations. In this case, the prior targets could have been at the same (S) or at a different (D) location as the target in the current trial. **d**, As in **b**, but for performance conditioned on the history of prior outcomes. In this case, the 1–4 trials preceding the current one could have been correct choices (C) or errors (E). Results in b–d are from the same simulation, which consisted of 100,000 trials. Black lines indicate 95% confidence intervals.

Although this particular task is hypothetical, it captures key elements of real tasks used to characterize visuomotor performance and its underlying neural mechanisms. Behavior in this task was simulated with a simple model (Methods) that generated many choices, one trial after another, such that selection history effects would be observed based on target location, target features, and outcome (correct/incorrect) — dependencies found in many prior studies [49, 50, 51, 52, 53, 54]. The underlying mechanisms whereby the model produces such sequential effects are most important, but to appreciate them we must first describe how those history effects manifest in the data.

To reveal the influence of selection history, we parsed the simulated choices (100,000 trials) according to their preceding histories as done in real psychophysical experiments (Fig 7b–d). In the case of target color, for instance, we computed the fraction of choices that were correct given that the color of the oddball in the previous trial was the same as in the current trial (Fig 7b, case S), or the fraction of choices that were correct given that the color of the oddball in the previous 3 trials was different from that in the current trial (Fig 7b, case DDD), and so forth. In the simulated data, performance got progressively better as the target color was repeated (Fig 7b, red bars going from 0.62 for S to 0.79 for SSSS), but got progressively worse as more color repetitions were followed by a switch in color (Fig 7b, blue bars going from 0.39 for D to 0.31 for DDDD). Analogous sorting of the simulated trials but now according to target location demonstrated a similar trend: the fraction correct increased noticeably when the target appeared repeatedly at the same location (Fig 7c, red bars going from 0.68 for S to 0.86 for SSSS); in contrast, performance dropped considerably when the prior target location was different from that in the current trial (Fig 7c, case D), and a location switch that followed up to 4 location repeats produced somewhat worse performance (Fig 7c, blue bars going from 0.45 for D to 0.38 for DDDD). Finally, the performance of the model also varied according to outcome history: choice accuracy was higher in trials that followed a correct choice than in trials that followed an error (Fig 7d, compare cases C and E), but the difference was relatively modest (0.54 vs. 0.47) and further repetitions had a barely visible impact (,:S 0.01) in either direction.

Based on these descriptive results, it is clear that the hypothetical subject’s performance on the current trial depends on the color of the prior targets, the location of the prior targets, and to a lesser degree, whether those prior trials were successful or not. But what drives these dependencies? And, are the underlying mechanisms responsible for these three effects independent or interrelated? These are questions that would be asked if the data were from a real experiment, and conditional independence can be used to address them.

The above analyses produced three sets of probabilities associated with the three individual selection histories. To understand the dependencies between the three variables (color, location, outcome), we can compute the probability of a correct choice given any given pairwise combination of such history terms and compare it to the probability predicted by assuming conditional independence. For example, consider the probability of success given that the oddball color in the prior trial was the same as in the current trial

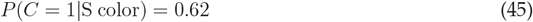

and the probability of success given that the prior choice was correct

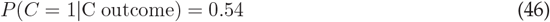

which were already obtained (Fig 7b, d). Then compute the probability of success given the joint condition

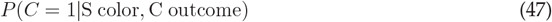

in two ways, (1) as measured from the data, by considering all the trials in which the preceding trial was both correct and of the same target color, and (2) as predicted by inserting the values in Equations 45 and 46 into Equation 15. The resulting numbers are 0.80 and 0.65, respectively (Fig 8a, top row, point marked SC). The difference between them is indicative of an interdependence between color and outcome. However, a more complete picture is obtained by repeating the same process for all the possible combinations of color and outcome going back one trial (Fig 8a, top row, points marked DE, DC, SE, SC). The resulting four points are clearly off the diagonal, and the actual, measured probabilities differ from the predicted ones in a particular pattern: the former deviate more from the overall probability of success, *P* (*C* = 1) = 0.51, which is simply the prior for a correct choice. That is, knowledge of the true joint history is more informative — it produces a more certain prediction — than expected based on conditional independence.

**Fig 8.**
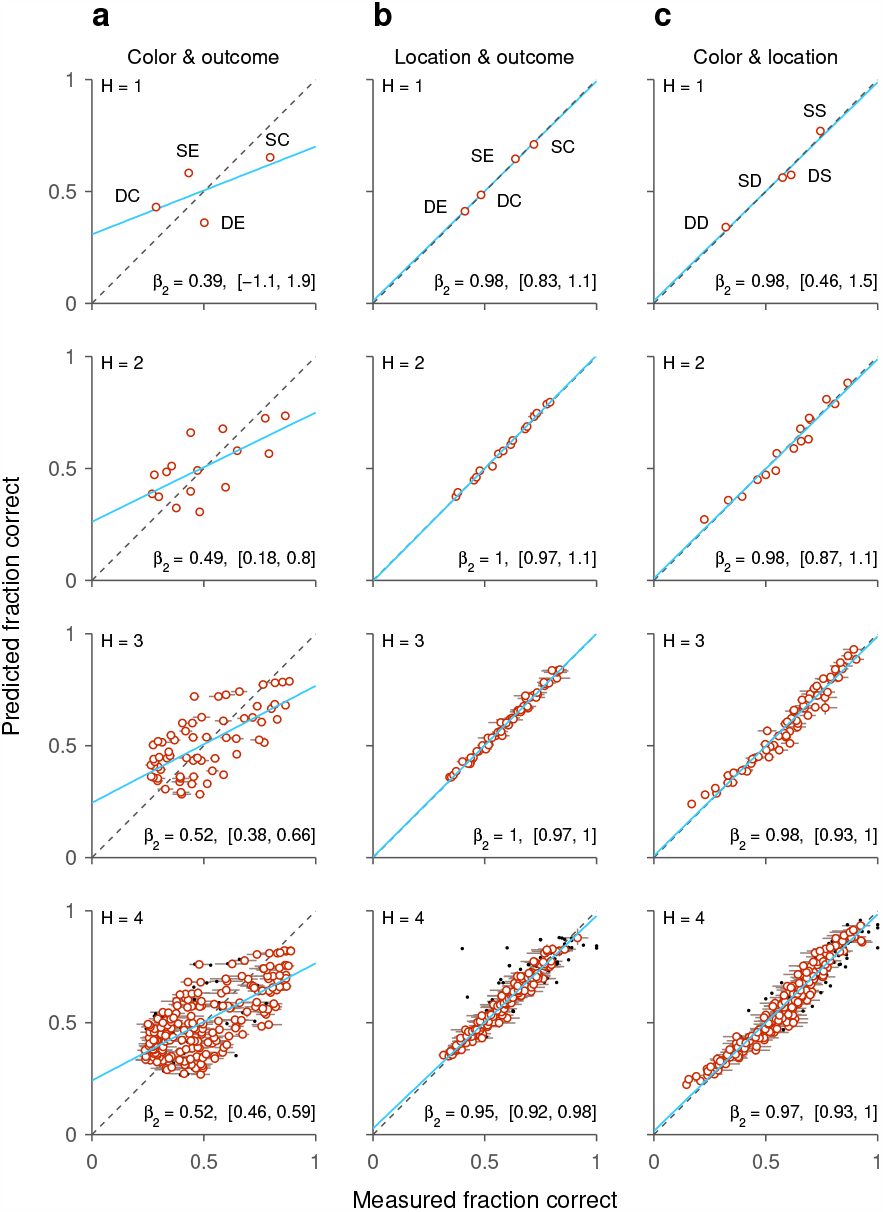
Interactions between selection history variables manifest as deviations from the predictions based on conditional independence. Each point indicates the probability of a correct choice given a joint set of histories, either measured from appropriately sorted, simulated trials (x axes) or as predicted by assuming conditional independence between the individual histories (y axes). **a**, Probabilities based on joint consideration of target color and choice outcome in prior trials. Rows correspond to histories going back 1–4 trials (*H* = 1, *H* = 2, *H* = 3, *H* = 4, respectively). Lines behind circles indicate 95% confidence intervals based on binomial statistics. Black dots are data points that had large uncertainties (95% confidence interval span *>* 0.15). Dashed diagonal lines mark equality between x- and y-axis values. Blue lines indicate linear regressions based on red circles. Regression slopes are indicated (*β*_2_, with 95% confidence intervals). **b**, Probabilities based on joint consideration of target location and choice outcome in prior trials. **c**, Probabilities based on joint consideration of target color and location in prior trials.

This pattern becomes more evident when the same comparison is made for joint histories going back 2, 3, and 4 trials (Fig 8, a, rows 2–4), which give rise to 4^2^, 4^3^, and 4^4^ different conditional probability terms, respectively. For example, the term

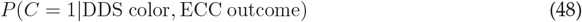

gives the probability of sucess given that the target colors of the three preceding trials were different, different, and the same as that in the current trial, and that the outcomes of those same preceding trials were error, correct, and correct. In all cases, the regression lines (Fig 8a, blue lines) for the resulting clouds of points cross the diagonal near the value of the prior (0.51) and have similar slopes *<* 1, confirming that the color and outcome histories are far from being conditionally independent. This suggests that there is some form of interaction between target color and outcome in the underlying mechanism whereby they influence the current choice.

A very different result is obtained when the same analysis is performed by combining target location and outcome histories (Fig 8b). Now the data points line up tightly along the diagonal, indicating high consistency between the empirical and predicted success probabilities. For histories going back 1–3 trials, most deviations from the diagonal are within the 95% confidence interval expected simply from the finite number of trials sampled in each condition; it is only when histories go back four trials that large deviations can be observed, typically for points that have large uncertainties (Fig 8b, black dots in bottom row indicate 95% confidence interval span *>* 0.15). Overall, the data suggest that target location and outcome do not interact; they influence each choice independently.

Based on their individual effects, it would appear that color and target histories exert qualitatively similar influences on task performance (Fig 7a, b). However, this superficial similarity is deceiving. Analyses of their effects jointly with outcome — when benchmarked against the performace expected based on conditional independence — reveals a stark difference between them (Fig 8a vs. b). And indeed, within the dynamics of the model, color and location histories act in entirely different ways mechanistically.

The model is very simple (Methods). It produces two kinds of response, guesses and informed choices. (1) During guesses, the model picks one of the four possible response locations based on where the oddball has previously appeared. Each of the four locations has a bias, and the probability of guessing toward location *j* is directly proportional to its bias. The biases are updated after every trial; whenever the oddball is presented at location *j*, the bias in favor of location *j* is increased and others are decreased. (2) During informed choices, there is a 0.95 probability of making a correct response toward the oddball and a 0.05 probability of making an erroneous response elsewhere (a lapse). The key to the model is that the probability of making an informed choice versus a guess depends on the match between the current color of the oddball and the prior colors experienced recently. Each of the two oddball colors (red and green) has a sensitivity. In each trial, the probability of making an informed choice depends on the sensitivity associated with the color of the current target: the probability is high if the oddball is red and the sensitivity for red is high, which is typically the case when prior targets have been red; and conversely, the probability is low if the oddball is red and the sensitivity for red is low, which is typically the case when prior targets have been green. Similarly when the oddball is green. Crucially, the sensitivities are reinforced after correct responses only. After a correct choice with a red oddball, the sensitivity to red is increased and that for green is decreased. Similarly for green. After an incorrect choice, both sensitivities decay slightly toward their neutral states.

In summary, the model simulates a subject that implements a simple strategy: if target and distracters can be discriminated (when sensitivity is high), the choice is informed and has a 95% chance of being correct, but if target and distracters cannot be discriminated (when sensitivity is low), the response is a guess that is biased toward recent target locations. Choice outcome enters into the dynamics, albeit indirectly; its only role is to gate the update of the color sensitivities. This is why the apparent effect of outcome history alone is weak and short-lived (Fig 7d).

So, color and location histories derive from fundamentally different mechanisms for driving performance, and this is highly consistent with the qualitative differences observed when success probability is conditioned on their respective joint histories combined with outcome (Fig 8a vs. b). Interestingly, further indication of their distinct mechanistic roles is obtained by a similar analysis in which success probability is conditioned on the joint history of color and location (Fig 8c). In this case, the data points also line up along the diagonal, indicating that, overall, color and location histories exert independent influences on performance.

In conclusion, this example illustrates the use of conditional independence as a benchmark for gaining some insight about how history variables modulate visuomotor performance. Specifically, analysis of the synthetic data suggested that the effect of color history was mediated by outcome (or reward, in a typical experiment), whereas that of location history was not, and furthermore, it indicated that the color and location influences were largely independent. All this was indeed in agreement with the model implementation. In additional computational experiments, alterations to the model produced corresponding alterations in the pattern of deviations from conditional independence, but the interpretation of the latter was always consistent with the former. The broader lesson is that, applied in this way, conditional independence may be a useful tool for assessing the presence of functional interactions between stochastic variables that modulate visuomotor performance.

## Conclusion

In this report we discussed the concept of conditional independence and its application as a statistical tool for characterizing a variety of evidence integration processes. The main result solves an elementary problem about predictability. Specifically, we showed that when two variables, *A* and *B*, are conditionally independent with respect to a third one, *C*, it becomes feasible to predict the state of the latter based on observation of the former using a simple computational recipe (Equation 16) and limited data, namely *P* (*C*|*A*), *P* (*C*|*B*), and *P* (*C*). We consider one last, intuitive example in order to highlight the simplicity of the problem and the usefulness of the solution as a benchmark for characterizing functional interactions between observable events.

Suppose the event of interest is whether your favorite baseball team wins (*C* = 1) or loses (*C* = 0) their next game. The team will be playing in Atlanta (*A* = 1), and their best player will be absent (*B* = 0). Based on their recent performance during the season, we know three things: the overall probability of a win, *P* (*C*), the probability of a win in Atlanta versus other cities, *P* (*C*|*A*), and the probability of a win with versus without their best player, *P* (*C*|*B*). Although the particular circumstance they face, playing in Atlanta without their best player, has not occured before, can we estimate the outcome with the information at hand? The answer is yes — if the game location and the best player exert independent influences on team performance, which is what conditional independence entails in this case.

Based on the standard definition of conditional independence (Equation 6), it may not be immediately clear why such a constraint would be interpreted as reflecting “independent influences” on the outcome variable *C*. The definition goes in the reverse direction; it is a statement about how knowledge of *C* affects the relationship between *A* and *B*. In terms of predictability, however, the two views are consistent. To see this, consider what independence means in practice.

Two events *A* and *B* are unconditionally independent (Equation 8) when their combined occurrence, *P* (*A, B*), can be accurately predicted based on limited knowledge, *P* (*A*) and *P* (*B*). If the actual combined (joint) probability is measured to be significantly smaller or larger than the expectation, *P* (*A*) *P* (*B*), then there is some sort of interaction or connection between the events. In the baseball example, *P* (*A, B*) corresponds to the probability that a game is in Atlanta (or elsewhere) and that the best player participates in it (or does not). Say the player is prone to miss games in Atlanta much more often than elsewhere because his favorite restaurant is there, and every time he visits he gets spicy food and invariably falls sick. Whatever the underlying reason, such an interaction would prevent us from calculating the probability *P* (*A, B*) by the usual approximation, *P* (*A*) *P* (*B*). Conversely, if the joint event is predictable, it means that the player is equally likely to be sick or injured in any city, and there is no interaction between events. Independence implies predictability and lack of interaction.

In the case of conditional independence the situation is entirely parallel. Now the two relationships *P* (*C*|*A*) and *P* (*C*|*B*) represent the separate statistical effects of events *A* and *B* on the outcome event *C*, and the probability *P* (*C*|*A, B*) represents the combined effect of *A* and *B* on *C*. If this probability can be accurately predicted based on limited knowledge, i.e., using *P* (*C*|*A*), *P* (*C*|*B*), and the prior *P* (*C*) only, then we should consider the two effects to act independently on *C*. If, on the other hand, the actual combined probability is measured to be significantly smaller or larger than the expectation, then there is some sort of interaction between the two effects. In the baseball example, say the best player is particularly crucial against Atlanta, so the probability of a win against Atlanta plummets to near zero when he is absent from the team. This interaction prevents us from predicting the probability *P* (*C*|*A, B*) by the appropriate approximation (Equation 16). Conversely, if there is no such interaction, if the impact of the best player is the same when playing against Atlanta as against other teams, then the approximation is valid and the result predictable. Thus, conditional independence implies predictability and lack of interaction.

Seen from this perspective, the definition of conditional independence (Equation 6) can be taken to mean that, when the state of *C* is known, the joint probability of events *A* and *B* becomes predictable based on limited information. As such, our results simply reveal a symmetry: when the condition is satisified, knowledge of *A* and *B* makes the state of *C* predictable based on limited information as well.

It is well established that conditional independence is a useful statistical tool for understanding how multiple sources of evidence are, or can be, integrated [1, 9, 10, 14, 38]. What we have emphasized here is its relevance for predicting outcomes conditioned on prior knowledge. In two examples, diagnosing a disease and inferring biological age, we demonstrated application of the analytical results to situations in which the primary goal would be to generate predictions. In two additional examples, characterizing the effectiveness of multisensory integration and determining how selection histories modulate visuomotor performance, the predictions served as benchmarks for characterizing (simulated) empirical data. The chosen examples are neither exhaustive nor highly detailed; they are meant as proofs of concept, and as guidelines for future applications to real data.

## Acknowledgments

We thank Denise Anderson for logistical and administrative support.

